# Human red blood cells express the RNA sensor TLR7 and bind viral RNA

**DOI:** 10.1101/2022.01.01.474694

**Authors:** LK Metthew Lam, Rebecca L. Clements, Kaitlyn A. Eckart, Ariel R. Weisman, Andy E. Vaughan, Nadir Yehya, Nuala J. Meyer, Kellie A. Jurado, Nilam S. Mangalmurti

## Abstract

Red blood cells (RBCs) express the nucleic acid-sensing toll-like receptor 9 (TLR9) and bind CpG-containing DNA. However, whether human RBCs express other nucleic acid-sensing TLRs and bind RNA is unknown. Here we show that human RBCs express the RNA sensor, TLR7. TLR7 is present on the red cell membrane and associates with the RBC membrane protein Band 3. RBCs bind synthetic single-stranded RNA and RNA from pathogenic single-stranded RNA viruses. RNA acquisition by RBCs is attenuated by recombinant TLR7 and inhibitory oligonucleotides. Thus, RBCs may represent a previously unrecognized reservoir for RNA, although how RNA-binding by RBCs modulates the immune response has yet to be elucidated. These findings add to the growing list of non-gas exchanging RBC immune functions.

**Significance:** Red blood cells interact with pathogens, cytokines, and CpG-containing DNA; however, their immune functions remain largely unexplored. Here, we report that RBCs can bind synthetic or viral RNA and express TLR7. TLR7 interacts with the RBC membrane protein Band 3 and this interaction is increased during acute infection with SARS-CoV-2. Our study suggests that RBCs may have immune functions mediated by viral RNA, and RNA-carrying RBCs can potentially be a reservoir for RNA-based diagnostics.

## Introduction

Despite lacking transcriptional and translational machinery, mature RBCs can exert immune function by binding to inflammatory mediators and pathogens using various receptors.^1-4^ Nucleic-acid sensing is an essential feature of the immune response, critical for triggering downstream inflammation. The nucleic acid-sensing TLRs (TLR3, 7, 8, and 9) are primarily localized intracellularly in the vesicular system.^5^ However, surface localization of these TLRs in some cell types may serve to facilitate cell-type-specific immune responses.^6-8^ We recently discovered that RBCs express surface TLR9 and bind CpG containing DNA, driving anemia and innate immune activation during inflammatory states; however, whether other TLRs are expressed on RBCs is unknown.^1,9^

Cell-free RNA (cfRNA), including microRNA, long non-coding RNA, tRNA, or fragments of mRNA, are present in circulation, often encased in vesicles.^10-14^ Although RNA is present throughout RBC development, mature RBCs are presumed to be devoid of RNA. Here we report that human RBCs are a reservoir of RNA, express the RNA sensor TLR7, and bind exogenous viral RNA.

## Results

Because we have previously identified the nucleic-acid sensing pattern recognition receptor (PRR) toll-like receptor 9 on erythrocytes and *Tlr7* is expressed in erythroid precursors, we asked if TLR7 would be present on RBCs.^15^ TLR7 is expressed on purified RBCs (**Fig. 1A)**. To ensure that the RBC preparation was devoid of platelets, which also expressed TLR7 and TLR9,^16,17^ we performed qRT-PCR for the platelet marker *Itga2b* (*CD41*) **(Fig. 1B)**. We did not detect *CD41* in our purified RBC samples, indicating that our RBC samples are free of platelet contamination. We detected TLR7 expression on RBCs from healthy donors and septic adult and pediatric patients with acute respiratory distress syndrome (ARDS) **(Fig. 1C-D)**. We confirmed our observation using confocal microscopy and two distinct clones of antibodies against TLR7 **(Fig. 1E)**. We next examined the interaction of TLR7 with the integral membrane protein Band 3 by proximity ligation assay (PLA) to confirm membrane localization of red cell TLR7 **(Fig. 2A-B)**. We found close interaction between TLR7 and Band 3, using two distinct antibody combinations **(Fig. 2A and C)**. Because we previously found that TLR9 and Band 3 interact in the RBC membrane, we asked if TLR7 interacts with TLR9.^1,9^ TLR7 interacts with TLR9 and Band 3 on the RBC membrane **(Fig. 2D)**. Collectively, these data establish the presence of the RNA-sensing pattern recognition receptor TLR7 on RBCs and suggests the presence of an immune sensing complex on the RBC membrane.

**Figure 1.**
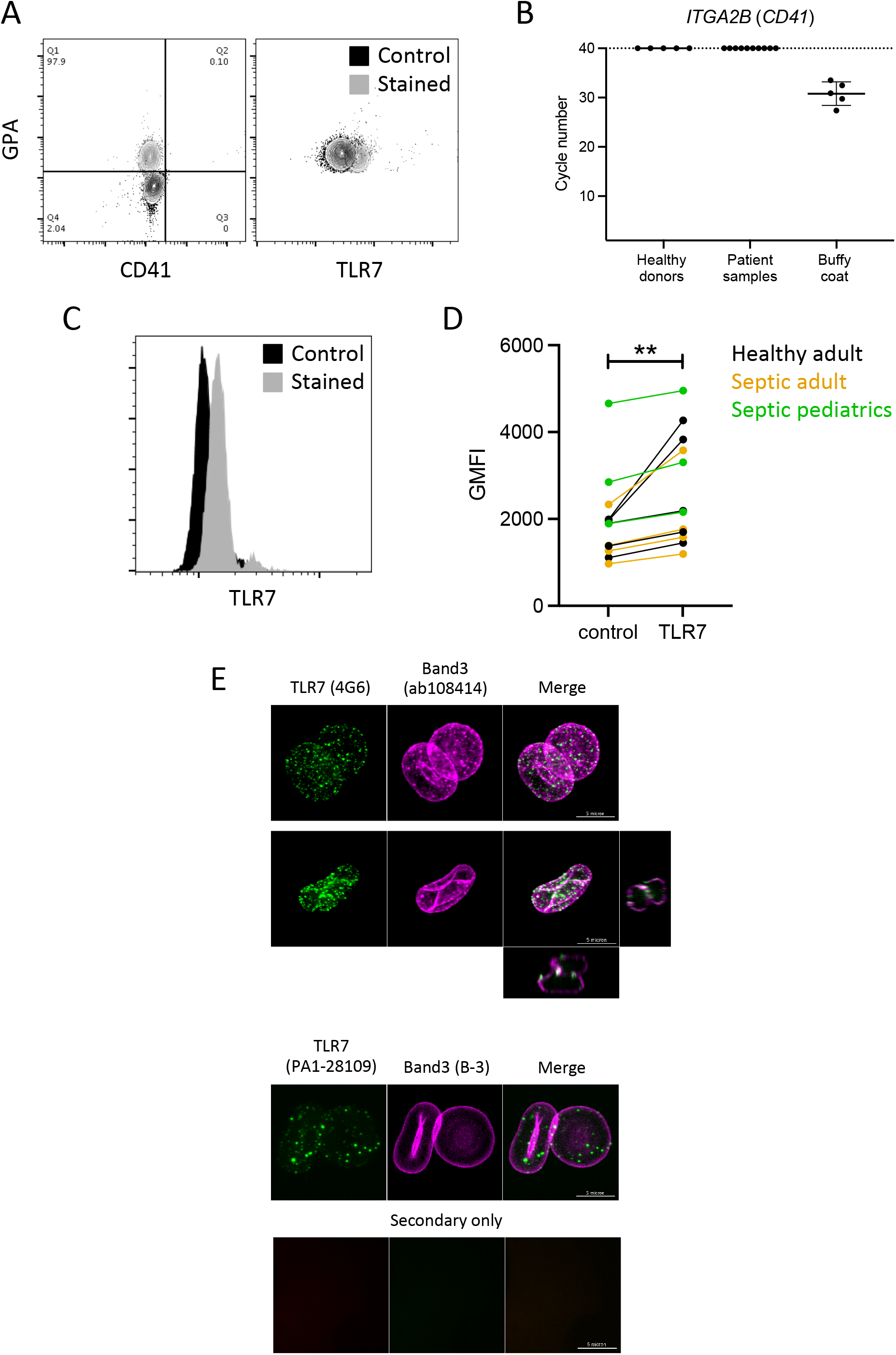
RBCs express TLR7. **(A)** Purified RBCs were stained for surface glycophorin A, CD41, and CD45, followed by TLR7 (4G6) staining after fixation and detergent treatment. **(B)** Relative levels of CD41 transcript in RBC preparations were quantified using qRT-PCR. Buffy coat was used as a positive control. **(C-D)** Flow cytometry detection of TLR7 on RBCs from healthy donors and septic patients. **(C)** Representative histograms of a healthy donor and **(D)** summarized data for geometric mean fluorescence intensity (GMFI). **(E)** Confocal micrograph for immunofluorescent staining of TLR7 and Band3 (RBC membrane) using two distinct pairs of antibodies. The clone of the antibody is indicated in parentheses. TLR7 staining in green, Band3 staining in magenta, and their corresponding merged images were shown. An orthogonal view is also shown for the 4G6-ab108414 antibody pair.

**Figure 2:**
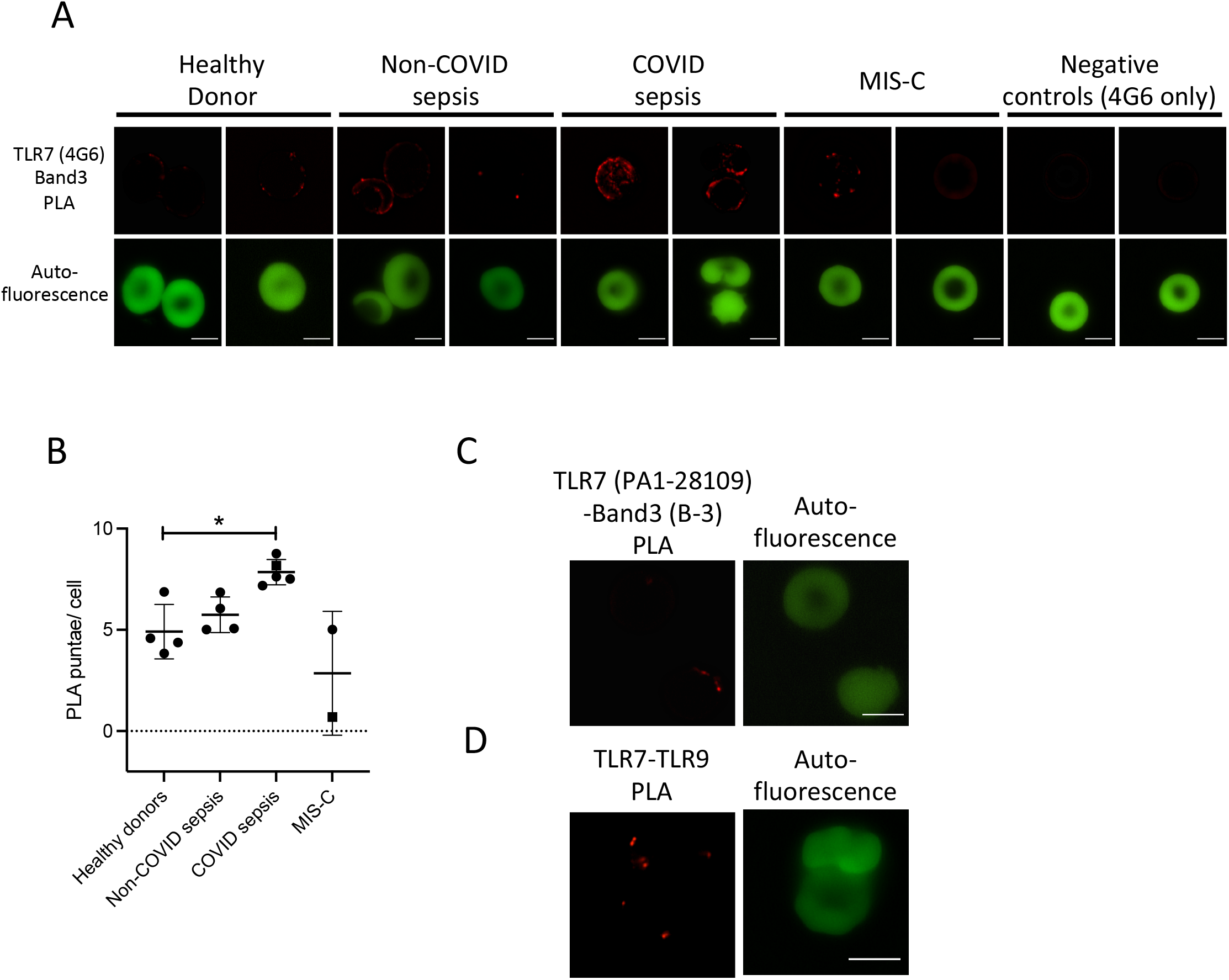
Interaction of TLR7 with Band3 and TLR9 on RBCs. **(A)** Proximity ligation assay (PLA) for TLR7 (clone 4G6) and band3 (clone ab108414) on RBCs from healthy donors and septic patients. Micrographs for PLA signal (red) and autofluorescence (green) of RBCs were shown. Each column of images represents RBCs from a distinct donor or patient. In negative control, only 4G6 antibody was used. Each pair of micrograph represents a unique donor or patient. **(B)** Quantification of PLA signals. Orange symbols depict COVID patients. Square (■) depicts pediatric patients. Difference between healthy donors and septic patients was evaluated by t-test, *: p<0.05. **(C)** PLA for TLR7 (clone PA1-28109) and Band3 (B-3) on RBCs from a healthy donor. **(D)** PLA for TLR7 (clone 4G6) and TLR9 (clone ab37154) on RBCs from a healthy donor. Scale bar represents 5μm.

We previously observed increased TLR9 expression on human RBCs during sepsis compared with healthy donors. However, in contrast to our observations for TLR9, we did not detect increased TLR7 on septic RBCs **(Fig. 1D)**. Because we observed Band 3-TLR7 interactions, and Band 3 alterations are reported during inflammatory states, including sepsis and malaria, we asked whether Band 3-TLR7 interactions would be altered during sepsis and COVID-19 associated sepsis.^18-20^ While TLR7-Band 3 interactions in RBCs from healthy donors were detected by distinct PLA punctae, RBCs from septic patients demonstrated a robust PLA signal with large punctae and aggregates of PLA punctae observed in bacterial sepsis and viral sepsis due to SARS-CoV-2 infection **(Fig. 2A-B)**. Because COVID-19 is associated with multi-system inflammatory syndrome (MIS-C), we also evaluated RBCs from two MIS-C patients, who were negative for active infection. We did not observe enhanced TLR7-Band3 interaction in the MIS-C patients. Thus, TLR7 association with RBC membrane proteins is increased during sepsis.

Because we detected TLR7 on RBCs, we examined whether RNA was present on RBCs under basal conditions. Using an RNA-selective dye, we measured the presence of RNA on RBCs **(Fig. S1A-B)**. Ribosomal RNA has previously been reported in RBCs.^21^ We, therefore, quantified 18S rRNA in RBCs from healthy donors and patients with sepsis; buffy coat preparations were used as a positive control. **(Fig. S1C-D)**. We next asked whether RBCs bind exogenous RNA. We found dose-dependent binding of single-stranded RNA oligoribonucleotide (ORN, RNA40) derived from the HIV-1 sequence **(Fig. 3A-B)**.^22^ Because the acquisition of DNA by RBCs led to morphological changes to the RBCs and masking of the anti-phagocytic epitope of CD47, we asked whether RNA binding would elicit the same response. RNA treatment of RBCs did not result in masking the CD47 anti-phagocytic epitope, suggesting that RBCs can distinguish RNA and DNA **(Fig. S2)**.

**Figure 3:**
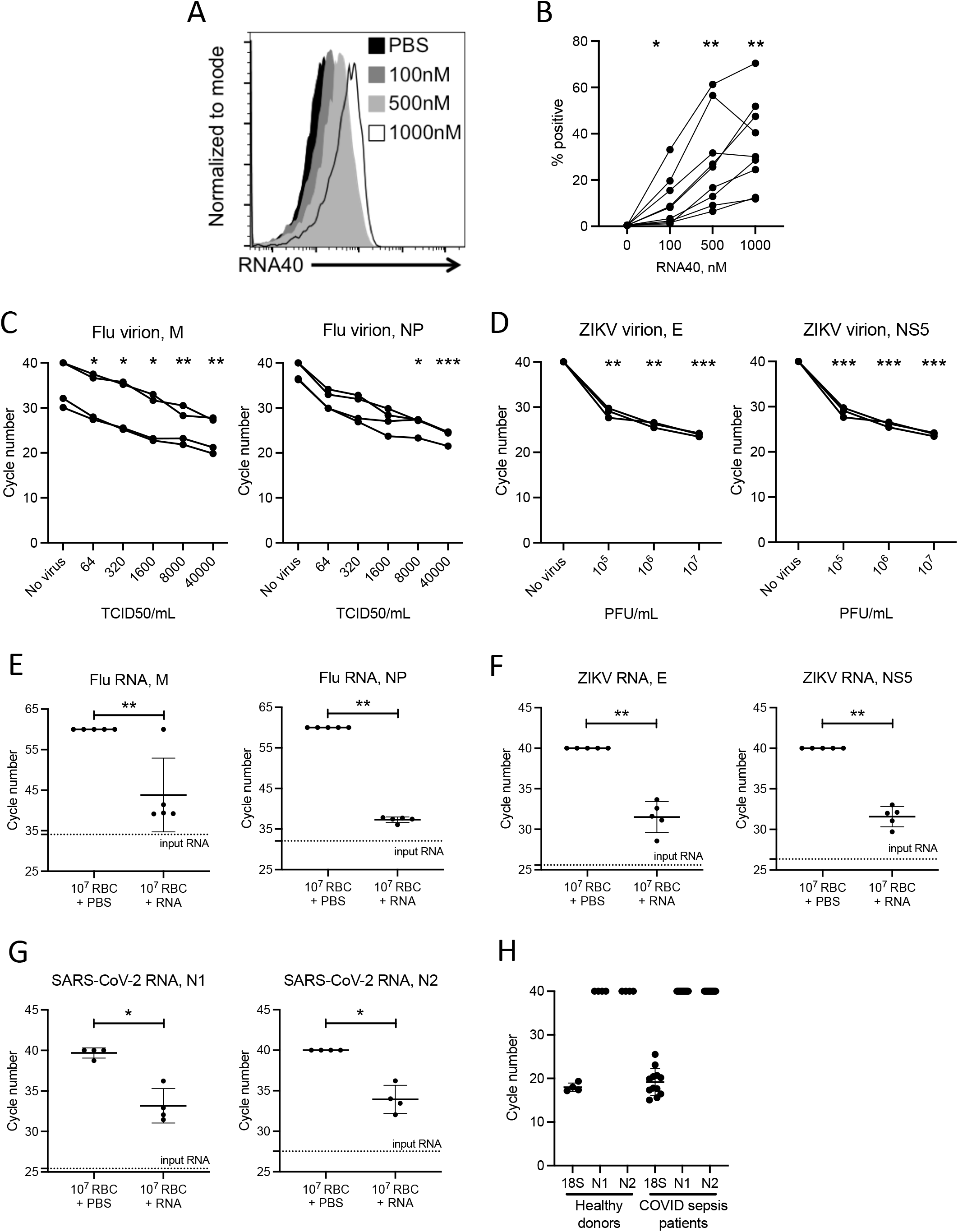
RBCs bind RNA. Binding of Cy5-labeled RNA40 oligoribonucleotide to RBCs from healthy donor. Representative histograms in **(A)** and summarized data in **(B)**. n=9, each line represent a unique donor. Differences in samples were evaluated with one-way ANOVA followed by Dunnett’s post-hoc test. **, p<0.01 **(C-D)** Binding of virus particles to RBCs. 10^7^ RBCs were incubated with indicated concentrations of influenza virus **(C)** or ZIKV **(D)** particles, and the relative amount of RBC-associated viral RNA was quantified with qRT-PCR for viral genes. Amplicons for matrix (M) and nucleoprotein (NP) were used for flu and envelope (E) and non-structural protein 5 (NS5) were used for ZIKV. Each line represents a unique donor. One-way ANOVA with Dunnett’s post-hoc compared to no virus. *: p<0.05; **: p<0.01; ***: p<0.001. **(E-G)** Viral RNA binding to RBCs. RBCs were incubated with 0.1ng influenza virus RNA **(E)**, 1ng ZIKV RNA **(F)**, or 1ng SARS-CoV-2 RNA. **(G)** and RBC-associated viral RNA was quantified with qRT-PCR. Two different amplicons for nucleocapsid (N1 and N2) were used for SARS-CoV-2. **(H)** RNA extracted from 5μL of packed RBC from healthy donor and septic COVID patients were quantified using qRT-PCR. 40 cycle is set as detection limit of the assay (n=12).

We next asked whether RBCs also acquire pathogen-derived RNA during infection **(Fig. S3A)**. Because influenza virus binds directly to red blood cells, we first asked whether viral RNA would be detectable on RBCs following incubation with influenza virus (strain A/Puerto Rico/8/1934 H1N1).^23^ Influenza virus RNA is detectable in RBCs following incubation with naïve RBCs **(Fig. 3C)**, indicating that RBCs acquire viral RNA following binding to virions. Zika Virus (ZIKV) is a mosquito-borne flavivirus that causes an acute febrile illness in adults and may persist in the circulation weeks after infection.^24^ It has recently been reported that ZIKV is detected primarily in the RBC fraction of transfusates, yet it is not known if ZIKV directly binds to RBCs or infects erythroid precursors.^25^ We found that naive RBCs bind ZIKV **(Fig. 3D)**. We next asked whether RBCs acquire viral RNA by directly binding to RNA; we incubated RBCs with RNA extracted from influenza A virions, Zika virions, or SARS-CoV-2-infected cells **(Fig. 3E-G, S3C-E)**. Although we observed heterogeneity in the ability of human RBCs to sequester viral RNA, viral RNA adhered to RBCs, suggesting that RBCs are capable of directly binding RNA from viruses.

Given recent reports of direct infection of erythroid progenitors with SARS-CoV-2 and our own findings of RBC-bound SARS-CoV-2 spike protein, we evaluated RBCs from patients hospitalized with COVID-19 **(Fig. 3H)**.^26,27^ We did not detect SARS-CoV-2 on RBCs from patients with COVID-19, although this is not surprising given that the day of blood sampling was on average ten days following symptom onset, and few papers have reported viremia in this population.^28,29^

We next asked whether blockade of ligand binding with TLR7-Fc or inhibitory ODNs would attenuate ssRNA acquisition by RBCs. RNA40 binding to RBCs was examined in the presence of increasing doses of soluble recombinant human TLR7-Fc. TLR7-Fc inhibits RNA40 binding in a dose-dependent manner, indicating that RNA40 is a substrate for TLR7 **(Fig. 4A-B)**. TLR7 is activated by polyuridine;^22^ we therefore tested if RBCs also bind to polyuridine ORN (15-mer ORN, polyU_15_). We found dose-dependent binding of polyU_15_ by RBCs; in addition, polyU_15_ competes with RNA40 for RBC binding **(Fig. 4C-D)**. We then asked if inhibitory oligonucleotides against TLR7 could inhibit RNA binding by RBC. ODN2088 and ODN20959 are CpG-containing oligonucleotides that inhibit TLR7 and TLR9 response in human and murine myeloid cells, whereas ODN105870 is derived from ODN20959 but contains a modified guanine that renders it inhibitory to TLR7 but not TLR9.^30,31^ We observed a dose-dependent inhibition of RNA40 binding by RBC by ODN2088 **(Fig. 4E-F)** and donor dependent heterogeneity in the inhibition of RNA binding by ODN2088. Similarly, donor heterogeneity was observed with ODN20959 and ODN105870 **(Fig. S4)**. We verified the ability of inhibitory ODNs to attenuate viral RNA acquisition by RBCs using viral RNA from ZIKV and SARS-CoV-2 **(Fig. 4G-H)**. Collectively these data suggest that synthetic and pathogen derived ssRNA binding to RBCs can be attenuated in the presence of TLR7 and TLR9 ligands.

**Figure 4:**
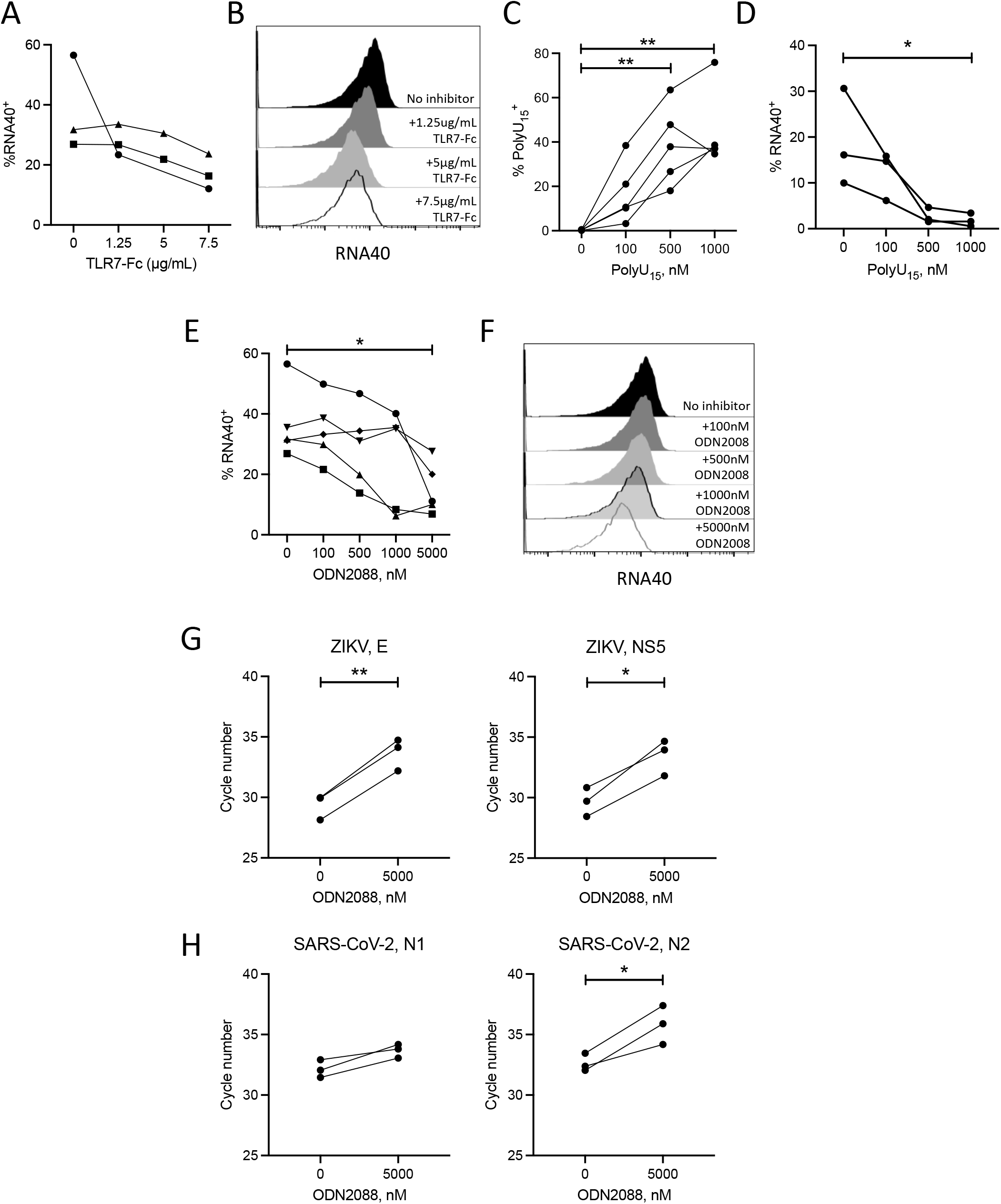
TLR inhibitors suppress RNA binding. **(A-B)** Binding of RNA40 to RBCs in the presence of TLR7-Fc. Percent RNA40^+^ cells **(A)** and a representative histogram **(B)** were shown. **(C)** Binding of polyU_15_ ORN to RBCs. **(D)** Competition of polyU_15_ and RNA40 ORNs. 500nM RNA40 were incubated with RBC in the presence of indicated amount of polyU_15_ ORN, and percent of cells positive for RNA40 was shown. **(E-F)** Binding of RNA40 to RBCs in the presence of ODN2088. Percent RNA^+^ cells **(E)** and a representative histogram **(F)** were shown. In **(A-E)**, differences across groups were performed with one-way ANOVA with Dunnett’s post-hoc comparing to untreated samples. Each line in A, C, D, and E represents a unique healthy donor. **(G-H)** Binding of ZIKV **(G)** or SARS-CoV-2 **(H)** viral RNA in the presence of 5000nM ODN2088. Differences between groups were compared using paired t-test. *: p<0.05; **: p<0.01

## Discussion

We demonstrate that TLR7 is expressed on RBCs and observe RNA binding by RBCs, which can be inhibited by TLR7 and 9 inhibitors. While these ODNs inhibit various endosomal TLRs responses, their binding activities are unclear. For example, ODN2088 competes with CpG-containing DNA for the C-terminus of TLR9 and inhibits downstream signaling, but how ODN2088 and other ODNs interact with TLR7 remains elusive.^32^ It is plausible that some of these ODNs only inhibit the downstream signaling but not the binding of TLR cognate substrates.

Because TLR7 and TLR9 are in close proximity on the RBC, it is also possible that ODN binding to TLR9 results in conformational changes that reduce the accessibility of ssRNA to TLR7. Further studies to examine this potential are underway. Moreover, it is unknown whether TLR7 expressed on RBCs retains its preference for GU-rich RNA. RBCs may also express alternative receptors for RNA, and it is known that RBCs express the purinergic receptors P2X7 and P2Y1.^33^

The reciprocal association of TLR7, TLR9, and Band 3 suggest a potential immune receptor complex on the RBC surface. ^5^ Band 3 is an anion channel expressed on the RBC surface that adopts multiple topologies and forms various multi-protein macro complexes to maintain optimal RBC structure and function. Clustering proteins of similar function in close proximity is reminiscent of lipid raft where molecules of related functions are concentrated to facilitate efficient signaling and regulation. In-depth investigation on the role of RBC TLR7 and its interacting partners is needed to establish the existence of such an immune sensing complex on the RBC membrane.

In our experiments, fixation and detergent treatment were needed for epitope accessibility of both TLR7 antibodies (4G6 and PA1-10826) that are raised against the TLR7 ectodomain; however, the proximal interactions of TLR7 and Band3 and susceptibility of RNA binding to TLR7 inhibitory oligonucleotides suggest TLR7 is localized on the RBC plasma membrane. Because the topology of Band 3 changes as RBCs age and during inflammatory conditions, it will be important to determine if TLR7 adopts non-canonical topology on the RBC membrane or if our findings are simply due to epitope accessibility.

Dating back to the HIV pandemic, transfusion-associated viral infection has long been a concern in blood banking. The advent of molecular screening and nucleic acid amplification techniques has dramatically reduced the risk of viral infection following blood transfusion.^34^ However, one lingering question has been the persistence of the virus in transfusates. Zika and WNV have both been reported to associate with RBCs in transfusates. West Nile virus (WNV) particles can be associated with RBCs,^35^ and persistent of RBC-associated ZIKV RNA has been reported in asymptomatic patients,^36^ yet whether the virus directly binds RBCs remained unknown. Consistent with previous reports, we observe the RBC binding of pathogenic viruses and RNA derived from them. Nucleic acids trapped in RBCs are reportedly nuclease resistant.^37^ This raises the question of whether RBC-associated RNA is protected from degradation by plasma RNases and is therefore detected following extensive storage even the host or donor no longer harbors the infectious virus. Because RBC-associated WNV retains infectivity, it is plausible that RBC-bound virions are shielded from the host immune response, although in vivo studies will be required to validate this hypothesis.

The ability of RBCs to capture exogenous RNA suggests that RBCs can serve as a reservoir for RNA in the circulation and may be exploited to develop diagnostics for infectious disease, cancer, or other inflammatory states where cell-free RNA may be abundant but not readily detectable in plasma samples. RBCs are associated with multiple types of RNA molecules, including miRNA and ncRNA,^38,39^ but the localization of these molecules is not well defined except the 7SL ncRNA, which is associated with cytoskeletal and membrane proteins in RBCs. Additionally, the function of these RNA molecules in the mature erythrocytes remains elusive. Our findings add to the growing literature demonstrating RNA association with RBCs.

Collectively, our data demonstrate that RBCs bind RNA and express the immune receptor TLR7. RBC-associated RNA may be valuable in disease diagnostics and monitoring. The discovery of immune receptors on RBCs suggests that RBCs possess unexplored immune function. Future work examining the characteristics of RBC-associated RNA during inflammation and the role of immune receptor expression on RBCs will address our knowledge gap on the long-overlooked non-gas exchanging immune function of RBCs.

## Materials and Method

### Pediatric Cohort

Subjects were selected from an ongoing prospective cohort study of intubated children with ARDS from the Children’s Hospital of Philadelphia CHOP). The study was approved by the CHOP Institutional Review Board, with informed consent obtained from caregivers. Specific inclusion criteria were: 1) acute (≤ 7 days of risk factor) respiratory failure requiring invasive ventilation, 2) age > 44 weeks corrected gestational age and < 17.5 years, 3) invasive ventilation via endotracheal tube, 4) <u>bilateral infiltrates</u> on chest radiograph, 5) oxygenation index ≥ 4 or oxygen saturation index ≥ 5 on <u>2 consecutive measurements at least 4 hours apart</u>; 6) invasively ventilated ≤ 7 days before meeting above radiographic and oxygenation criteria; 7) invasive blood drawing access (central venous catheter, arterial catheter, or blood-drawing IV). <u>Exclusion criteria</u> were 1) weight < 3 kilograms, 2) cyanotic congenital heart disease, 3) tracheostomy, 4) invasively ventilated for > 7 days when meeting criteria above, 5) cardiac failure as <u>predominant</u> cause of respiratory failure, 6) primary obstructive airway disease (asthma, bronchiolitis) by judgement of clinician as the primary cause of respiratory failure, 7) alternative known chronic lung disease as cause of respiratory failure (cystic fibrosis, eosinophilic pneumonia, interstitial pneumonitis, pulmonary hemosiderosis, cryptogenic organizing pneumonia), 8) severe moribund state not expected to survive > 72 hours, 9) any limitations of care at time of screening, or 10) previous enrollment in this study.

### Study approval

Studies involving human subjects were approved by the University of Pennsylvania Institutional Review Board. Healthy volunteers between ages of 18 and 65 years gave written informed consent prior to inclusion.

### Sepsis Cohort

RBCs were obtained from patients enrolled in the Molecular Epidemiology of Severe Sepsis in the ICU cohort (MESSI) or inpatient subjects positive for SARS-CoV-2 enrolled in the MESSI-COVID study at the University of Pennsylvania.^40,41^ Patients were eligible if they presented to the medical intensive care unit (MICU) with strongly suspected infection, at least 2 systemic inflammatory response syndrome (SIRS) criteria, and evidence of new end organ dysfunction in accordance with consensus definitions by the American College of Chest Physicians. Exclusion criteria included a lack of commitment to life-sustaining measures, primary reason for ICU admission unrelated to sepsis, admission from a long-term acute care hospital, previous enrollment, or lack of informed consent. Human subjects or their proxies provided informed consent.

### RBC isolation

To separate blood fractions, blood samples were centrifuged at 3000 g for 10 minutes. Plasma and buffy coat were saved or aspirated depending on the experiment. Red blood cells were purified from the remaining packed red cell fraction using magnetic-assisted cell sorting (MACS) or leukoreduction filters as previously described.^1,9^ MACS-isolated RBCs were frozen and used in qPCR or fixed in 0.05% glutaraldehyde for staining (see below) whereas leukoreduced RBCs were immediately used in all functional assays. For RBCs used for qRT-PCR, and 5μL of packed RBCs were frozen after centrifugation and removal of the buffy coat and plasma.

### Flow cytometry

For surface staining, 250,000 cells were washed with FACS buffer (PBS + 2% FBS) and blocked in anti-human Fc block for 30min on ice, followed by staining with CD45, CD41, and glycophorin A antibodies for 30min on ice. Antibody information is presented in Supplemental table 1. For TLR7 staining, cells were washed three times in PBS and fixed with 0.05% glutaraldehyde for 10 minutes at room temperature, followed by 3 washes with FACS buffer. Cells were permeabilized in 0.1% triton X-100 diluted in FACS buffer for 15 minutes, followed by 3 washes with FACS buffer. Cells were stained with 5μg FITC-conjugated anti-TLR7 antibody, 4G6, or isotype for 1hr in the dark. Cells were washed twice in FACS buffer prior to analysis with a flow cytometer (BD Fortessa).

### Immunofluorescence and microscopy

RBCs were fixed at 10^7^ cells/mL in 0.05% glutaraldehyde and permeabilized as above. RBCs were then blocked in PBST (PBS + 0.05% tween 20) supplemented with 1% BSA for 1hr at room temperature under gentle shaking. For each staining reaction, 10^6^ fixed and permeabilized RBCs were blocked in PBST supplemented with 1% BSA and 5% goat serum for 1hr at room temperature. Blocked RBCs were stained with primary antibodies listed in Supplemental table 1 diluted in PBST with 1% BSA overnight at 4°C. Cells were washed three times in PBST and stained in secondary antibodies raised in goat (Jackson ImmunoResearch) for 1hr at room temperature under dark, followed by 3 washes in PBST. Cells were resuspended in PBS and mounted on Fluoromount G. Confocal micrographs were acquired with VT-iSIM (Visitech).

### Transmission electron microscopy

RBCs were fixed in TEM fixative (4% Formaldehyde, 3.5% glutaraldehyde in 0.1M sodium cacodylate). Samples were processed and stained by Electron Microscopy Core at the University of Pennsylvania. The TLR7 antibody 4G6 was used in immunogold staining.

### Proximity ligation assay

10^6^ fixed and permeabilized RBCs were stained in polypropylene tubes (Falcon 352052). PLA (DuoLink, Sigma) were carried out according to the manufacturer’s protocol. After the final wash, cells were resuspended in no more than 10uL PBS and mounted on Fluoromount G. Images were acquired with a Nikon 2A microscope. At least 5 fields were imaged for each sample and these images were counted by two blinded personnel.

### RNA staining

SYTO RNAselect (SYTOR, Invitrogen) was used to stain RNA on RBCs according to the manufacturer’s protocol. Briefly, 500nM SYTOR diluted in PBS was incubated with 250,000 freshly isolated RBCs for 20min at room temperature under gentle shaking. Afterwards, cells were washed with PBS three times prior to analysis by flow cytometer (BD Fortessa).

### RNA extraction and reverse transcription

Previously frozen RBCs were thawed on ice. RNA was extracted using RNeasy Plus Kit (Qiagen) according to manufacture’s protocol. At the initial step, RBCs were lysed in 600μL lysis buffer and at the final step, RNA were eluted in 30μL RNase-free water. 8μL of freshly isolated RNA were reverse transcribed to cDNA using Superscript First Strand Synthesis system (ThermoFisher) following manufacture’s protocol.

### qRT-PCR

Quantitative RT-PCR was performed using the QuantStudio7 Flex system (Applied Biosystems) with 384 well plates. The amount of cDNA was quantified using PowerUp SYBR or Fast Taqman master mix following manufacturer’s protocol. The primers or Taqman assay mix were listed in Supplemental table 2. For quality controls, the resulting amplicons were resolved on 1.2% agarose gel in TBE buffer whenever necessary. Alternatively, for SYBR green-based assays, melting temperature was compared to controls.

### Oligoribonucleotide binding and inhibition

ORNs and ODNs used in this study were purchased from Integrated DNA technologies. In each binding reaction, 250,000 RBCs were resuspended in 100uL sterile PBS in polypropylene tubes. Fluorescently labeled (Cy5) ORNs (RNA40 and polyU_15_) were added to each reaction in volume no larger than 5uL. In assays where inhibitors were needed, 500nM RNA40 was titrated against various concentrations of inhibitors. Sequences for the used oligonucleotides were listed in Supplemental table 3. Recombinant human TLR7-Fc was obtained from R&D systems. Tubes were sealed in parafilm and incubated at 37°C for 2hr on a nutator in the dark. Afterwards, cells were washed with 1mL sterile PBS twice prior to analysis on a cytometer. For RNA competition with polyU_15_, 500nM fluorescein labeled (6-FAM) RNA40 was used.

### CD47 staining

RBCs incubated with RNA40 were washed with 1mL FACS buffer twice and probed with 5μg anti-CD47 antibody (CC2C6, Biolegend) for 1hr. Cells were washed with FACS for three times prior to analysis by a flow cytometer.

### Virion and viral RNA binding

Virus stocks and RNA were a kind gift from Dr. Andrew Vaughn (Influenza virus) and Dr. Kellie Jurado (Zika virus and SARS-CoV-2). All viral particle and RNA binding assays were performed at 100μL reaction volume in 2.0 mL round bottom DNA/RNA lo-bind tubes (Eppendorf, 86-924). All viruses and corresponding RNA were diluted in sterile PBS. 10^7^ RBCs were used in virus particle binding assays. For viral binding inhibition, 1ng viral RNA and 10^7^ RBCs were used. RBCs were incubated with virus particles or RNA for 2hr at 37°C on a nutator. Tubes were rotated at 1hr to ensure suspension of cells. For virus particle binding, RBCs were washed with 1mL PBS three times and frozen at -80°C until RNA extraction. For viral RNA binding assays, RNA-RBCs mix were overlaid on 500μL 30% sucrose in PBS at 4°C. Cells were pelleted by centrifugation at 13000g for 3min, followed by 2 additional washes in 1mL PBS and storage at -80°C until RNA extraction.

## Figure legends

**Figure S1: RNA is present in RBCs**.

**(A-B)** RBC-associated RNA was stained with SYTO RNA. Representative contour plot in **(A)** and summarized data for donors in **(B)** n=5, differences in samples were evaluated with paired t-test. *, p<0.05 **(C-D)** 18S rRNA detection in RBCs. Total RNA extracted from varying number of RBCs or buffy coat cells from healthy donors were quantified using qRT-PCR. **(C)** Amplicon of 18S from RBCs. **(D)** Comparison of relative level of 18s rRNA in buffy coat cells and RBC.

**Figure S2: RNA40 does not result in conformational change in CD47**

RBCs from healthy donor were incubated with RNA40. Afterwards, cells were washed and probed for CD47 expression with antibody CC2C6. **(A)** Representative contour plot and **(B)** summary of data were shown. n=3

**Figure S3: RBCs bind viral RNA**

**(A)** Schematics for the experiments to determine virus or viral RNA binding to RBCs. **(B)** Representative amplification curves for viral RNA binding. **(C-E)** 0.1ng of flu RNA **(C)**, 1ng of ZIKV RNA **(D)**, or 1ng SARS-CoV-2 **(E)** RNA were incubated with indicated number of RBCs. RBC-associated RNA were quantified.

**Figure S4: Inhibition of RNA binding by ODN20959 and ODN105870**

Binding of RNA40 to RBCs in the presence of ODN20959 (A) or ODN105870 (B). Percent RNA40^+^ cells were shown

