## Supplementary figures and images for "Human red blood cells express the RNA sensor TLR7 and bind viral RNA"

### supplemental figure 1-4

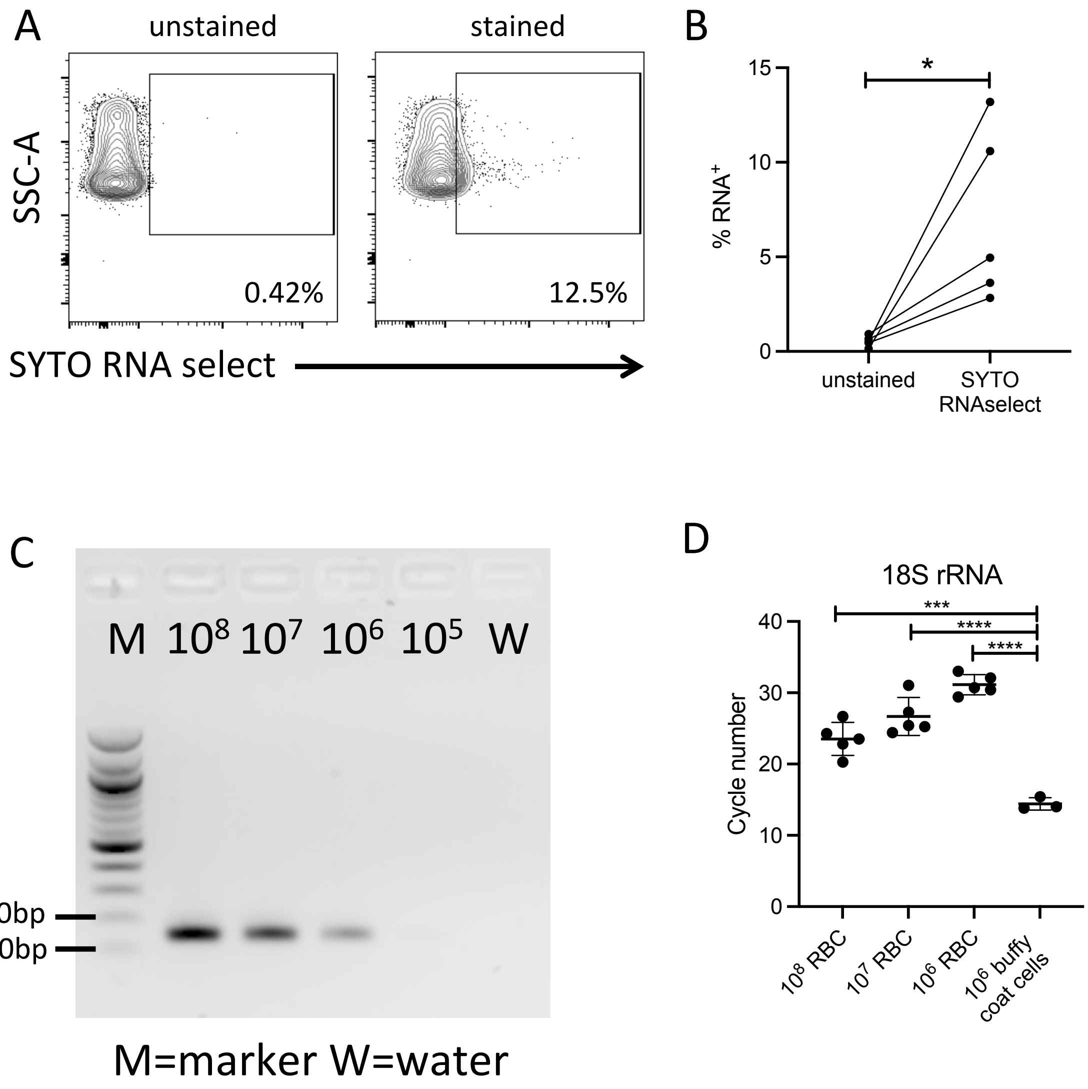

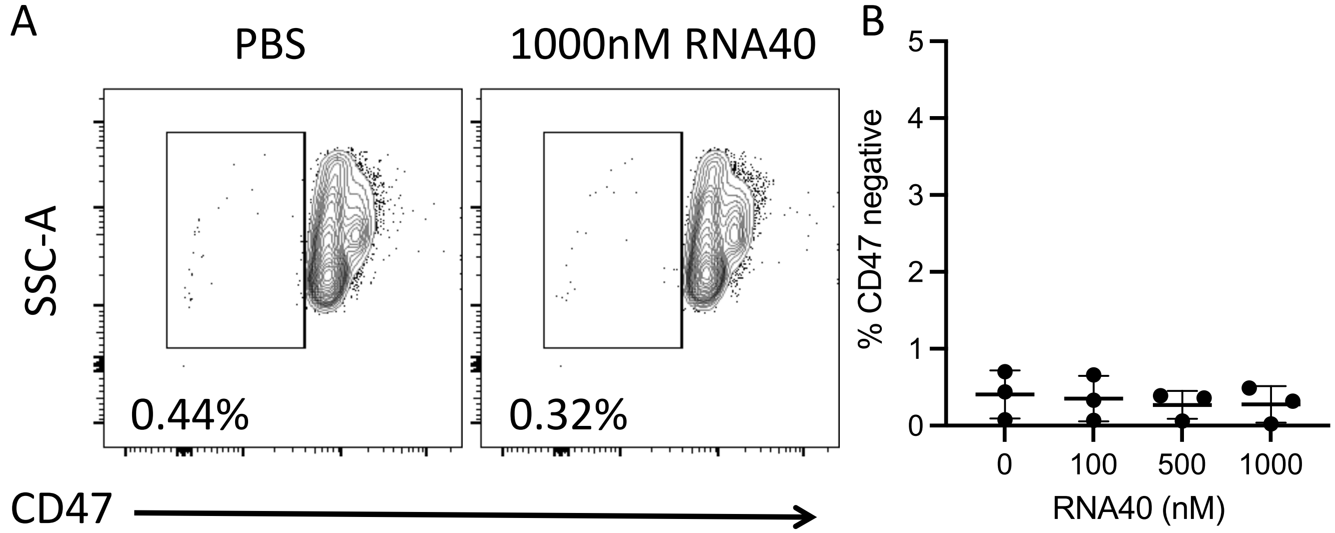

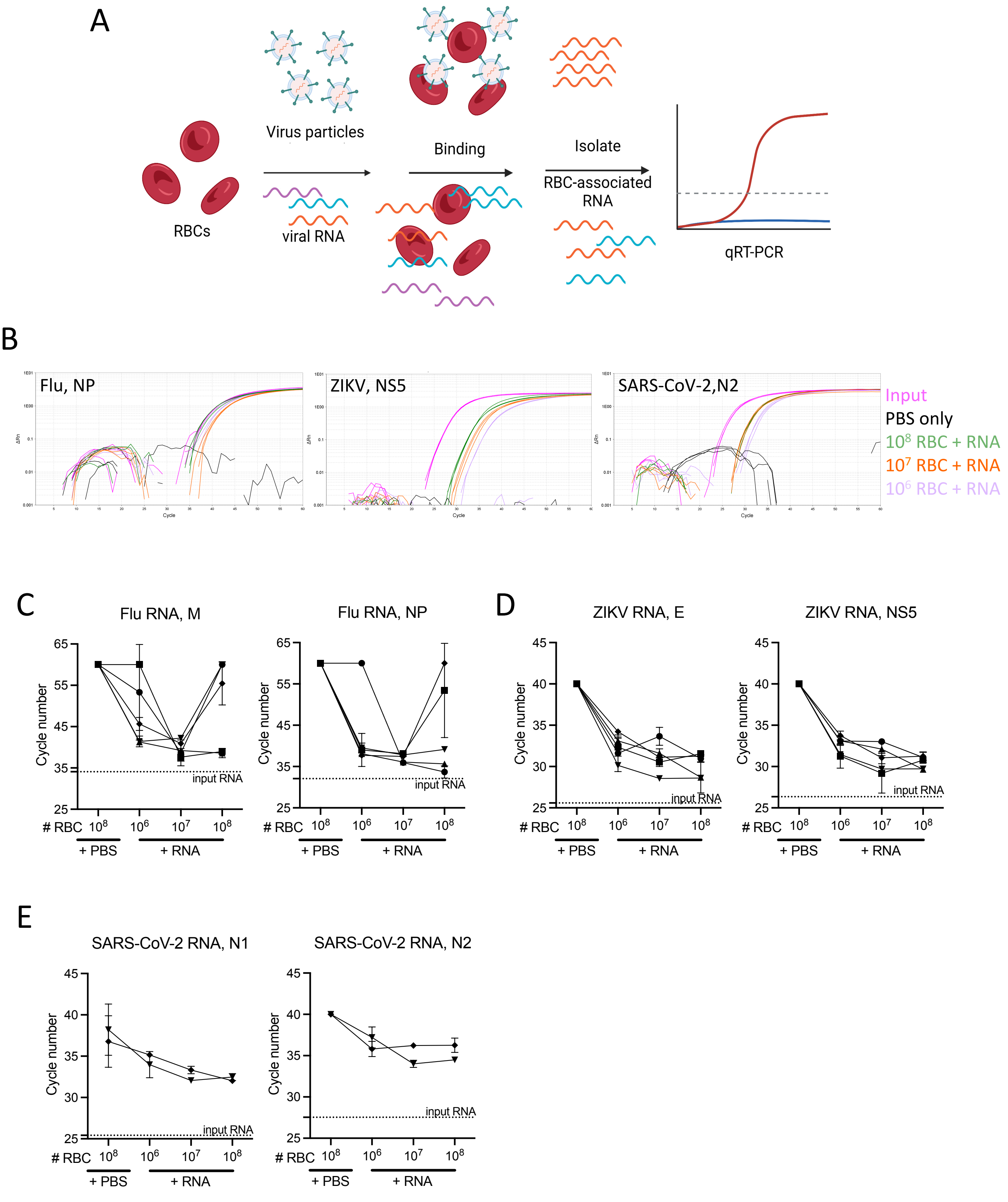

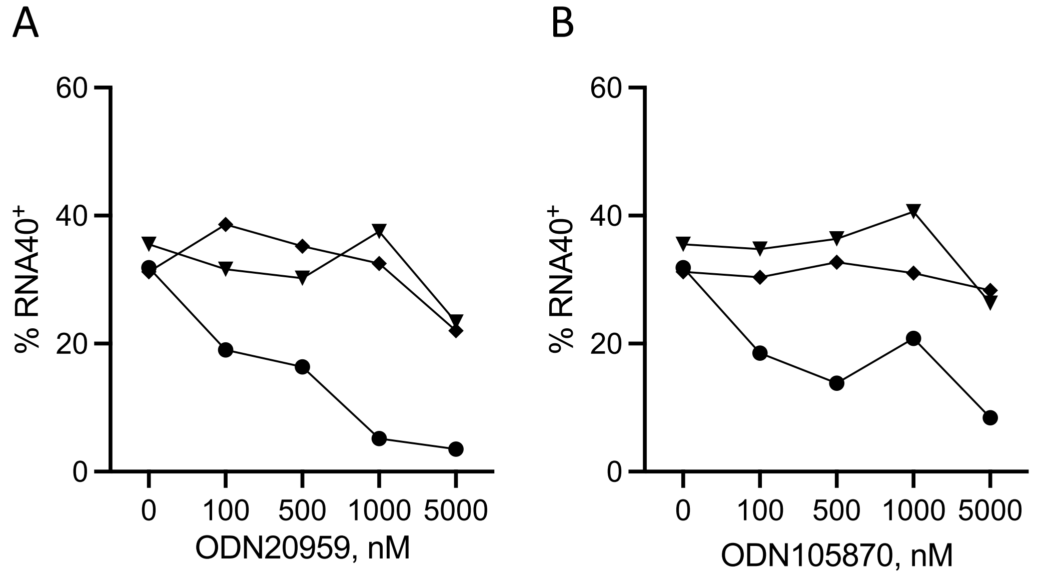
