## supplemental table 1-3 for "Human red blood cells express the RNA sensor TLR7 and bind viral RNA"

**Supplemental Table 1. Information on the antibodies used in this study**

| Antigen | Clone | Fluorophore | Usage | Vendor |
|---|---|---|---|---|
| TLR7 | 4G6 | FITC | FC: 5µg | Novus Biologicals |
| | | N/A | IFA, PLA: 2.5µg/mL | Novus Biologicals |
| TLR9 | PA1-28109 | N/A | IFA, PLA: 5µg/mL | Invitrogen |
| | ab37154 | N/A | IFA, PLA, 5µg/mL | Abcam |
| Band3 | ab108414 | N/A | IFA, PLA: 1.4µg/mL | Abcam |
| | A-3 | N/A | IFA, PLA, 1µg/mL | Santa Cruz |
| CD41 | HIP8 | APC | FC: 7.5µg/mL | Biolegend |
| GPA | HI264 | PE | FC: 0.5µg/mL | Biolegend |
| CD45 | 2D1 | AlexaFluor700 | FC: 5µg/mL | Biolegend |
| CD47 | CC2C6 | APC | FC: 5µL/assay | Biolegend |

FC: flow cytometry; IFA: Immunofluorescence; PLA: proximity ligation assay

**Supplemental Table 2. Gene expression assays used in this study[42,43]**

| Gene | Taqman assays/primers |
|---|---|
| CD41 | Taqman: Hs01116228_m1 |
| 18S | Taqman: Hs99999901_s1 |
| 28S | Taqman: Hs03654441_s1 |
| SARS-CoV-2 | 2019-nCoV RUO Kit, IDT |
| ZIKV-E | Forward: TTG GTC ATG ATA CTG CTG ATT GC |
| | Reverse: CCT TCC ACA AAG TCC CTA TTG C |
| ZIKV-NS5 | Forward: GGC CAC GAG TCT GTA CCA AA |
| | Reverse: AGC TTC ACT GCA GTC TTC C |
| Flu-PR8-M [40] | Forward: GGA CTG CAG CGT AGA CGC TT |
| | Reverse: CAT CCT GTT GTA TAT GAG GCC CAT |
| Flu-PR8-NP [41] | Forward: GAC GAT GCA ACG GCT GGT CTG |
| | Reverse: ACC ATT GTT CCA ACT CCT TT |

**Supplemental Table 3. Oligonucleotides used in this study**

| Oligonucleotide | Sequence |
|---|---|
| RNA40-Cy5 | rG*rC*rC*rC*rG*rU*rC*rU*rG*rU*rU*rG*rU*rG*rU*rG*rA*rC*rU*rC-Cy5 |
| RNA40-FAM | rG*rC*rC*rC*rG*rU*rC*rU*rG*rU*rU*rG*rU*rG*rU*rG*rA*rC*rU*rC-6-FAM |
| PolyU$_{15}$-Cy5 | rU*rU*rU*rU*rU*rU*rU*rU*rU*rU*rU*rU*rU*rU*rU-Cy5 |
| 2088 | T*C*C*T*G*G*C*G*G*G*G*A*A*G*T |
| 20959 | T*A*A*T*G*G*C*G*G*G*G*A*A*G*T |
| 105870 | T*A*A*T*G*G*C*E*G*G*G*A*A*G*T |

"*" denotes phosphorothioate bond; "r" denotes ribonucleotide; "E" denotes 7-deaza-2'-deoxyguanosine
